# The Maintenance of Sex and David Lack’s Principle

**DOI:** 10.1101/035832

**Authors:** Joachim L. Dagg

## Abstract

Combining George C. Williams’ idea that evolutionary constraints prevent asexual mutants from arising more frequently in low fecundity organisms, like mammals and birds, with an earlier one by David Lack that the brood size of these organisms has an optimum, and producing larger broods reduces their fitness, leads to a novel hypothesis about the maintenance of sex in them. All else equal, the eggs of an asexual mutant female should simply start developing without fertilisation, and there is no reason to assume that they would stop doing so after the optimal number of offspring has been produced. Without a way to control their reproductive output, asexual mutants should over-reproduce and suffer a cost of doing so. Experimental studies suggest that the cost of enlarged broods could limit the advantage of asexual mutants considerably. Moreover, research discovered that increased reproductive effort reduces immune functions of low fecundity organisms. This offers a surprising synthesis between Williams’ constraint and Hamilton’s parasite hypothesis on maintaining sex in low fecundity organisms: Compromised immune functions of asexual hosts may render them susceptible rather than adaptation on the side of parasites to overcome host resistance.

## INTRODUCTION

George C. Williams was instrumental in raising attention to the fact that the evolutionary maintenance of sexual reproduction was an unsolved problem. Many animal and plant species reproduce sexually and half the offspring is male, yet the males contribute nothing but genes to reproduction. If mutants in such a population could produce as many offspring asexually as the sexual females produced daughters plus sons, on average, their growth rate would be twice as large. Whereas half the offspring of sexual females would be non-contributing males, all the offspring of a mutant would contribute to growth rate. It has therefore often been called the ‘twofold cost of sex.’ Williams (1975) conceived this as the cost of producing offspring that is less related (r = 0.5) than clonal offspring (r = 1)*—cost of meiosis*, whereas Maynard Smith (1978) conceived it as the cost of investing half the reproductive resources into non-contributing males*—cost of males*.

The advantage gained by asexual mutants would only be twofold, however, if they produced as many offspring as sexual individuals with an even sex allocation. Conversely, a mutant would still gain some advantage as long as it produced more than half as many offspring. Nevertheless, sexual reproduction persists in most species and is not lost evolutionarily. Williams (1975: 7) called this the ‘paradox of sex.’

### Williams’ life history models

Williams (1975: 15) likened sexual reproduction to buying different lottery tickets and asexual reproduction to buying twice that number of copies of the same ticket, instead, and he (Williams 1975: 17) assumed that siblings compete among each other. Although evidence shows that clonal offspring is genetically variable (e.g., Lushai et al. 2003; Monti et al. 2012), the assumption that sexual offspring will be more variable than asexual may still hold.

Williams also suggested life-cycle types (verbal models) that should maintain sexual reproduction. All his models worked by sexual reproduction generating a variety of genotypes, selection leaving only a tiny minority of genotypes that are the very fittest for the environmental conditions experienced, but the probability of these conditions recurring for their offspring being negligible (Williams 1975: 14, 16, 59). Genotype fitness will be highly heritable within local habitats, but not between them (Williams 1975: 4, table 1).

**The aphid-rotifer model** (Williams 1975: chap. 2) supposes organisms living in discontinuous habitat patches, like ephemeral hosts or temporary puddles, that alternate between asexual and sexual reproduction. These patches are exploited by producing clones, because the heritability of fitness within a patch is high, but each patch is transient. Conversely, asexual reproduction will be useless for colonising new patches, because the patches vary in such a way that the fittest genotype in one patch will not survive in the next (Williams 1975: 17, 20). The dispersing offspring is therefore produced sexually (Williams 1975: 23; see also Bell 1982: 110f).

**The strawberry-coral model** (Williams 1975: chap. 3; see also Bell 1982: 112) supposes sessile organisms in continuous habitats. While a clone could theoretically spread without meeting a boundary confining the habitat, it would soon meet conditions limiting its further spread (Williams 1975: 26). With competition from neighbours the spread of a clone would be limited even more (Williams 1975: 27). As genotype fitness is not heritable between localities, producing dispersal stages asexually would be futile under the strawberry-coral model, again, because Williams (1975: 32f) assumed that the environmental gradients make it unlikely that a clone’s fitness will be higher in any locality other than the one it already occupies. (Cf. Combosch & Vollmer (2013) on an exceptional coral that can also produce dispersing larvae asexually.)

**The elm-oyster model** (Williams 1975: chap. 4) is similar to the strawberry-coral model with the difference that the local competition between genotypes is best run by growing large somata rather than many clonal ramets. In elms, for example, a single vertically growing stem might be better at shading out competitors than lateral ramets (Williams 1975: 42). The asexual mode of reproduction has consequently been reduced to zero in organisms of the elm-oyster type (see also Bell 1982: 113f). Williams (1975: 35) regarded a life-cycle with no asexual mode as an extreme case of both modes in varying degrees. The reproductive output of mature individuals is supposedly of the same order as that of a whole clone of the other life-cycle types (Williams 1975: 43).

The **other models** proposed by Williams (1975: chap. 5) suppose mobile organisms with high fecundity. In the triton model (Williams 1975: 50ff), the adults do not clone themselves or grow large somata, in order to exploit a local habitat, because they are mobile. Their movement, although considerable on a local scale, is negligible in comparison with that of their larvae, which are dispersed for month by oceanic currents. Furthermore, the adult can actively search out favourable conditions and avoid unfavourable ones, whereas the larvae are passively carried along and meet uncertain conditions. In the cod-starfish model (Williams 1975: 52ff), widely dispersed propagules colonise a neighbourhood that permits only one unpredictable phenotype to survive. Sexual reproduction will, again, more likely produce this winning genotype than asexual reproduction.

### Low fecundity organisms

Although Williams (1975) managed to come up with life history arguments that could explain the maintenance of sex in various organisms with high fecundity, he failed to come up with a model life-cycle that could explain it in obligately sexual organisms with low fecundity such as mammals, birds and many insects (presuming even sex ratio and males that contribute nothing to reproduction beyond mating). Williams (1975:62) defined low fecundity as less than 10^3^ offspring per average genotype. For these, he appealed to historical constraints preserving sexual reproduction, when it has ceased to be adaptive (Williams 1975: 44 and chap. 9, see also Bell 1982: 89). He assumed that asexual would replace sexual reproduction as soon as the constraints were overcome:

> “If and when any form of asexual reproduction becomes feasible in higher vertebrates, it completely replaces sexual. So in these forms sexuality is a maladaptive feature, dating from a piscine or even protochordate ancestor, for which they lack the preadaptations for ridding themselves.” (Williams 1975: 103)

Hamilton (1975: 179; 1980: 288; et al. 1990: 3566, 3572; 2001: 17, 605f) took this sub-problem of maintaining sex in obligately sexual organisms with low fecundity as his point of departure and benchmark of success. He did not think that constraints could properly explain the maintenance of sex in these organisms:

> “Williams himself seems to have despaired of showing advantages for sexuality for low fecundity organisms and concludes, in effect, that most practise sex because they haven’t found suitable tricks for eliminating it yet (can he really believe this for so many vertebrates?).” Hamilton (1975: 179, see also Seger & Hamilton 1988: 178)

Hamilton arrived at the conclusion that recombination protects against quickly adapting parasites in general. His red-queen model explains the maintenance of sex in high as well as low fecundity organisms with genotypes that are either sexual or asexual.

> “We simulated a host population of 200 individuals that are either sexual hermaphroditic or else all-female and parthenogenetic; the difference is controlled by a single gene.” Hamilton et al. (1990: 3566)

(Jokela et al. (2003), Salathé et al. (2008), Lively (2010) and Vergara et al. (2014: Introduction) review theory and evidence on this hypothesis.)

Williams (2000), however, pointed out that organisms of the strawberry-coral type disappoint expectations from the red-queen theory. With irony about ‘authorities,’ he indicated that the asexual offspring, here runners, stay close to their parents and hence their parasites, whereas the sexual propagules disperse.

> “No real scientists ever agree on everything, and Bill and I had a brief conflict last year at the Stony Brook conference. I am not convinced that adaptation by local pathogens to parental genotypes need be the major problem solved by sexuality. I think that the general unpredictability of offspring environments is what provides the main advantage. This issue is most appropriately settled not by modelling or data gathering but by consulting authorities. For a reliable insight on the significance of sexuality there are many appropriate authorities, but one that is especially clear is the strawberry plant (Fragaria). Offspring that develop immediately in the parents’ environment, with pathogens adapted to those parents’ genotypes, will not be sexually produced; whereas those that develop at variable times in the future, over a large range of habitats will be. The allocation of resources to sexual and asexual reproduction must be that which balances the two-fold cost of meiosis by the advantage of genetic diversity among widely dispersed seeds.” Williams (2000)

If coevolving pathogens were the force maintaining sexual reproduction in hosts, the dispersing offspring should be produced asexually and the staying (philopatric) offspring sexually. The opposite is true in the strawberry’s exemplary life-cycle.

Indeed, Hamilton never modelled a species with two genotypes, both using both modes of reproduction in their life-cycle, where one genotype produces dispersing offspring sexually and philopatric offspring asexually and the other does vice verse. Lively (2010) reviewed red-queen models, all pitting obligate sexual and asexual genotypes against each other—not genotypes using both modes in different situations. This, however, only challenges the universality of the red-queen theory. (See Brown et al. (1995); Hanley et al. (1995); Vernon et al. (1996); Weeks (1996); Ben-Ami & Heller (2005; 2008); Tobler & Schlupp (2005); Killick et al. (2008); Kotusz et al. (2014) for evidence on exceptions to red-queen predictions.)

### The cost of reproduction as a possible constraint

Let us return to Williams’ initial reasoning. He thought that the absence of asexual mutants from low fecundity organisms was a matter of constraints (Williams 1975: 44, chap. 9). No practical biologist interested in the maintenance of sexual reproduction would be lead to work out the detailed consequences experienced by asexual mutants he considered lacking in the first place. In fact, asexual low fecundity organisms are rare rather than lacking (for cases of asexuality in vertebrates see Vrijenhoek et al. 1989; Booth et al. 2014: supporting information). Yet what else should we do if we wished to understand why they are rare? Where Neiman (2004) and West-Eberhard (2005) already worked out physiological consequences, Dagg (2006) sociobiological ones, Kono (2006), Lushai & Loxdale (2007), Gorelick & Carpione (2009) epigenetic ones, and Mogie (2013) a cytological one, the following proposes a life-history constraint.

All else equal, the eggs of a mutant female should simply start developing without fertilisation, and there is no reason to assume that they would necessarily stop doing so after they have produced their optimal offspring number. Consider a mammalian mutant female, whose eggs developed parthenogenetically. Every oestrus an egg would ovulate from the ovaries and start developing in the oviduct without fertilisation. Unlike unfertilised eggs, such embryos would produce the required hormones and, on reaching the uterus, implant instead of degenerate. If nothing intervened with this development, they should produce more offspring than optimal for their fitness.

At this point, a look at David Lack’s principle can be instructive. He suggested that natural selection should maximise the number of viable offspring at the end of the period of parental care.

> “as litter size increases, the proportionate mortality among the young increases, and that eventually an optimum litter-size is reached, beyond which the initial advantage of an extra youngster is more than offset by the rise in proportionate mortality.” Lack (1948:46)

Williams (1966) refined this hypothesis as a trade-off between current reproductive effort and remaining reproductive value (cost of reproduction). There should be an optimal brood size beyond which producing more offspring will reduce rather than increase the individual’s lifetime reproductive success. This theory gets complicated further through environmental factors and individual condition rendering the cost of reproduction variable in time and space as well as individually (e.g., Boyce & Perrins 1987; Grant et al. 2000; Chambert et al. 2013).

Experimental studies corroborate the general idea that brood size has an optimum and reproducing more reduces fitness (Lindén & Møller 1989; Dijkstra et al. 1990; Koskela 1998; Aubret et al. 2003; Koivula et al. 2003; Santos & Nakagawa 2012; Aloise King et al. 2013; Thomson et al. 2014; Dugas et al. 2015), though the cost can be paid primarily by offspring, mothers, or fathers (in species with contributing males). Though a representative review of the relevant literature is beyond the scope of this article, the following examples illustrate the cost of enlarged brood size in two vertebrate species. Dijkstra et al. (1990) experimentally enlarged the brood size of kestrels by two chicks and offspring mortality rose from 1.5% in control groups to 19.4% in enlarged broods. Aubret et al. (2003) enlarged the clutches of ball pythons from 6.50 to 9.75 eggs on average. Where only 3.1% of the eggs failed to hatch in the control group, 25.6% died in the enlarged clutches. While this does not suffice to annul the hypothetical twofold advantage of asexuality, it shows the potential of this cost to constrain asexual mutants whose advantage is already lower (< 20%) for other reasons.

Moreover, research (e.g., Siva-Jothy et al. 1998; Fedorka et al. 2004; Hanssen et al. 2005; Harshman & Zera 2007; Christe et al. 2015; Treanor et al. 2015; Reedy et al. in press) discovered a link between reproductive effort and immune dysfunction. This offers a surprising synthesis between the constraint and the parasite hypothesis. Over-reproduction may compromise immune functions of asexual low fecundity organisms. If that renders them susceptible, infection will be a matter of chance rather than pathogen adaptation to overcome resistance.

Both the cost of reproduction and the immune function in asexual low fecundity organisms can be tested using know cases of asexuality in them (see Vrijenhoek et al. 1989; Booth et al. 2014).

